# Dopamine Accumulation Dynamics Control Time Perception

**DOI:** 10.64898/2026.01.13.699158

**Authors:** Tiezhan Lu, Junyou Sun, Yiping Lu, Ang Li, Zilong Gao, Wenzhi Sun

**Affiliations:** Academy for Advanced Interdisciplinary Studies, Peking University; Beijing 100871, China; Chinese Institute for Brain Research; Beijing 102206, China; Beijing Institute for Brain Research, Chinese Academy of Medical Sciences & Peking Union Medical College; Beijing102206, China; School of Basic Medical Sciences, Capital Medical University; Beijing 100069, China; Department of Psychiatry, First Hospital/First Clinical Medical College of Shanxi Medical University; Taiyuan, 030001, China; Shanxi Key Laboratory of Artificial Intelligence Assisted Diagnosis and Treatment for Mental Disorder, First Hospital of Shanxi Medical University; Taiyuan, 030001, China

**Keywords:** Time Perception, Ventral Tegmental Area, Dopamine Accumulation, Dopamine Clock Hypothesis, Internal Clock Hypothesis

## Abstract

Accurate timing is fundamental to perception and behavior, and the dopaminergic system plays a key role. According to the dopamine clock hypothesis, elevated dopamine accelerates the interval clock, while reduced levels slow it. However, optogenetic activation of substantia nigra pars compacta (SNc) dopaminergic neurons paradoxically disrupts this model, suggesting greater complexity. To resolve this paradox, we monitored ventral tegmental area (VTA) dopamine activity in mice performing a time perception task. We observed a ramping-to-threshold pattern, where slower ramping corresponded to longer durations. Optogenetic activation accelerated time estimation, whereas inhibition prolonged it. Systemic dopamine elevation had no effect, but reuptake inhibition accelerated timing. Strikingly, prolonged high dopamine levels slowed time estimation, correlating with slower accumulation due to increased reuptake. Our findings challenge the dopamine clock hypothesis, demonstrating that time perception depends not on absolute dopamine levels but on their temporal accumulation dynamics. This revised framework reconciles previous contradictions and highlights the role of dopamine accumulation rates in temporal cognition.

## INTRODUCTION

Accurate timing is essential throughout life, forming the foundation of temporal awareness that, enables event prediction, action coordination, and the understanding of past and future events. Various brain regions are crucial for time perception, including the prefrontal cortex (1–3), sensory cortex (4–6), motor cortex (7–9), hippocampus (10–13), and notably, the midbrain dopamine system (14–16). Evidence from early lesion studies (17–19) and dopamine-related disorders such as Parkinson’s disease (20–22), schizophrenia (23), and ADHD (24, 25) suggests that dopamine deficiency impairs time perception.

Pharmacological manipulations targeting dopamine have shown that increasing dopamine levels speeds up the internal clock, whereas reducing dopamine levels slows it down (26, 27). This phenomenon is known as the dopamine clock hypothesis (28–30). Advances in genetic manipulation have enabled specific modulation of dopamine pathways. For example, dopamine transporter (DAT) knockout either eliminates time perception or speeds up the internal clock (31), whereas D2 receptor overexpression slows it down (32). However, direct optogenetic manipulation of substantia nigra pars compacta (SNc) dopaminergic neurons produced results contradicting the dopamine clock hypothesis (15), suggesting that dopamine regulates time perception through mechanisms beyond simple activity levels.

To resolve these contradictions and elucidate the mechanisms underlying dopaminergic regulation of time perception, we developed a time perception task in which mice needed to wait for a delay before receiving a reward (33), allowing us to probe dopamine’s role in time perception. Using fiber photometry, we recorded activity in ventral tegmental area (VTA) dopaminergic neurons and identified a ramping-to-threshold dopaminergic activity pattern during the waiting period. Notably, the ramping rate was slower for longer durations, suggesting a direct relationship between dopaminergic activity dynamics and time estimation. Furthermore, transient optogenetic activation of VTA dopamine neurons during the waiting period accelerated time estimation, whereas inhibition slowed it, consistent with prior pharmacological findings.

However, pharmacological interventions revealed greater complexity. Systemic elevation of dopaminergic activity with morphine did not alter time estimation, whereas dopamine reuptake inhibition (with AHN) significantly accelerated it. Further experiments combining optogenetic stimulation of VTA DA neurons and a dopamine-specific fluorescent sensor in the nucleus accumbens (NAc) demonstrated that prolonged high dopamine levels slowed time estimation. This effect was accompanied by slower dopamine accumulation with higher reuptake efficiency, suggesting a dynamic interplay between dopaminergic activity and its temporal effects. Together, these findings challenge the notion that time perception is directly determined by the magnitude of dopaminergic activity. Instead, our results highlight the critical importance of dopamine accumulation dynamics during timing, offering a revised framework for understanding dopamine’s role in time perception.

## RESULTS

### Mice learned to wait accurately for 2s or 3s to receive rewards

We first trained mice to perform a time perception task to earn rewards, dividing the training into two phases. In phase one, the water-restricted mice engaged in a one-arm foraging task as previously described (33). The time spent in the waiting zone was designated as the waiting duration, while the time taken to move from the waiting zone to the water reward was termed the running duration. After 7 days of training, both the average waiting duration and running duration significantly decreased (p < 0.0001, Fig. S1C to E). All mice adapted their strategies, effectively reducing both waiting and running times to obtain rewards more efficiently.

Next, the mice were trained using a time perception paradigm. They were divided into two groups: one required to wait longer than 2 seconds and the other longer than 3 seconds to receive rewards. After 18 days of training, the waiting durations for both groups increased significantly and stabilized, while the coefficient of variation (CV) of waiting duration gradually decreased (Fig. 1C to F & Fig. S1G to J). In the 2-second group, 29.8% of waiting durations were less than 2 seconds (Fig. 1D), while 30% of the 3-second group waited less than 3 seconds (Fig. 1H). Running times did not show significant differences throughout the training (Fig. S1G to J). These findings indicate that the mice successfully learned to wait for the target durations of 2 or 3 seconds, demonstrating stable behavioral performance.

**Fig. 1.**
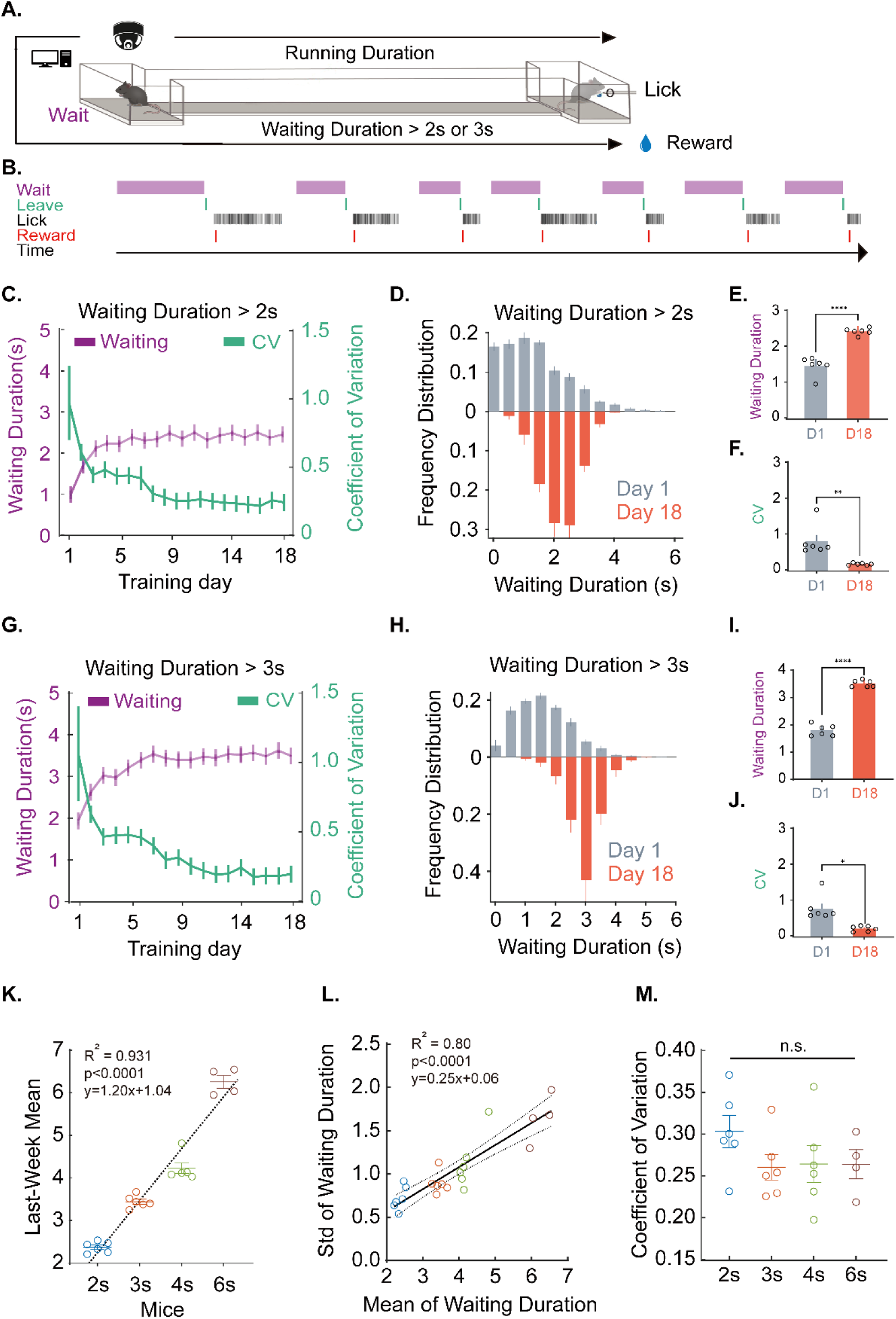
Scalable training of timed waiting behavior. (A) The Schematic of the behavioral apparatus. (B) Training protocol and task logic. (C–F) Outcomes for the 2-second threshold cohort (n=6): waiting durations increased from ∼1.5 s to ∼2.5 s and stabilized (Days 1–18), with the coefficient of variation (CV) declining from 0.8 to 0.2; on Day 18, <28% of trials were below 2 s. (G–J) The 3-second cohort (n=6) showed a similar trend, with mean waiting duration increasing from ∼1.8 s to ∼3.4 s and CV decreasing from 0.89 to 0.23; <30% of trials on Day 18 were below 3 s. (K–M) Scalability analysis across cohorts (2s, 3s, 4s, 6s) revealed a linear relationship between group means and standard deviations, with no significant difference in CV between cohorts. Data points represent: Blue: 2-second cohort (n = 6); Orange: 3-second cohort (n = 6); Light green: 4-second cohort (n = 6); Brown: 6-second cohort (n = 4). Data are presented as mean ± SEM. Statistical significance was determined by paired Student’s t-test (*p < 0.05; p < 0.01;p < 0.001;***p < 0.0001).

In many time perception tasks, behavioral performance often aligns with the scalar property of time perception (34). To investigate this further, we trained additional groups of mice to wait for 4 seconds and 6 seconds. During the final week, we analyzed the waiting durations across all four groups. The mean waiting duration increased linearly with the target duration across these groups (Fig. 1K, R² = 0.931, p<0.0001), indicating temporal accuracy. Additionally, the mean waiting durations were linearly correlated with the standard deviation of waiting duration among different mice and groups (Fig. 1L, R² = 0.80, p<0.0001). The CV of waiting duration did not differ significantly across the four groups (Fig. 1M), consistent with scalar variability reflecting Weber’s Law in time perception. These results demonstrate that the mice learned to accurately time intervals for rewards.

### VTA dopaminergic neurons ramped to threshold with varying slopes during time perception

To investigate the activity of VTA dopaminergic (DAergic) neurons during time perception, we recorded their calcium signals throughout the task (Fig. 2A). The raw calcium signal revealed a gradual ramping of DAergic activity after the mice entered the waiting zone, peaking as they exited (Fig. 2B). To analyze the dynamics of this ramping activity, we categorized the calcium signals based on waiting duration from the final week of training (Fig. 2C and G). Ramping activity was detected in 78.5% of trials for the 2-second group and 80.1% for the 3-second group (Fig. S2C and F). We further examined the waiting times associated with ramping versus non-ramping activities (Fig. S2B and E). In the 2-second group, the average waiting time for ramping trials was 2.5 seconds, compared to 3.6 seconds for non-ramping trials (t = 4.131, p < 0.01). In the 3-second group, ramping trials averaged 3.5 seconds of waiting, while non-ramping trials averaged 4.2 seconds (t = 4.738, p < 0.01). In both groups, non-ramping waiting times were significantly longer.

**Fig. 2.**
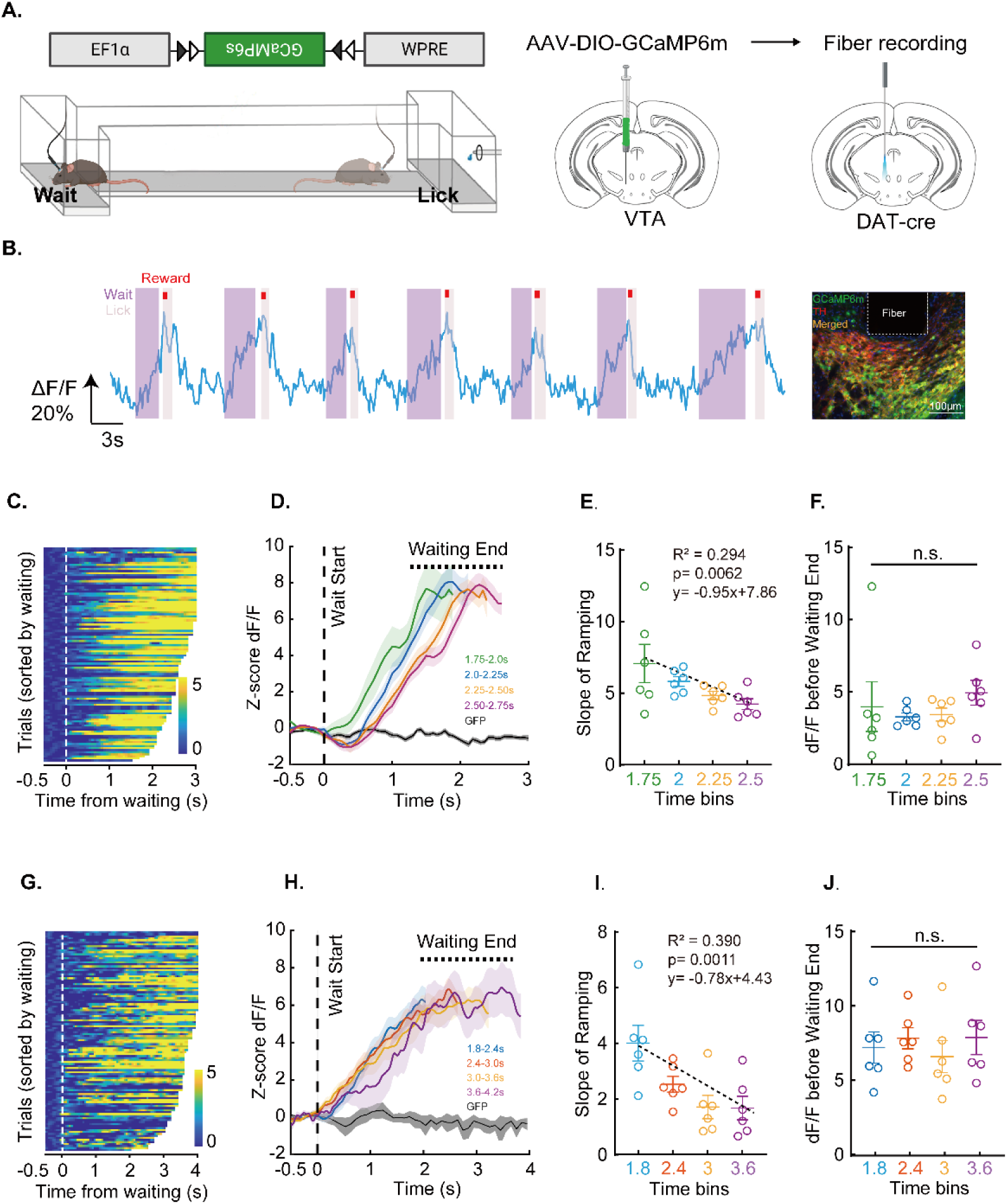
Slope analysis of ramping VTA-DA neuronal calcium signals during waiting. (A) Schematic of GCaMP6 expression in VTA of DAT-cre mice and fiber photometry setup. (B) Temporal alignment of behavioral events and calcium signals: Lavender: Waiting window. Pink: Licking. Red: Water reward (10 μL). Blue: Raw calcium trace. (C, G) Heatmaps of calcium signals sorted by waiting duration for representative mice in the 2-second (C) and 3-second (G) threshold cohorts. (D, H) Average z-scored ΔF/F curves during waiting, plotted for specific time bins relative to wait-end, alongside GFP control (gray), for the 2s (D) and 3s (H) cohorts. Signals ramped to a similar threshold at wait-end across bins. (E, I) Linear regression revealed that mean ramping slopes negatively correlated with waiting duration in both the 2s (E; R²=0.294, p<0.01) and 3s (I; R²=0.390, p=0.0010) cohorts. (F, J) Mean z-scored ΔF/F immediately before waiting end showed no significant difference between time bins in either cohort (ANOVA). Statistical annotations: ∗p<0.05, ∗∗p<0.01, ∗∗∗p<0.001, ∗∗∗∗p<0.0001.

We categorized trials into distinct groups based on waiting duration (ranges: 1.2 to 2.8 seconds for the 2-second group; 1.8 to 4.2 seconds for the 3-second group). Average signal traces indicated that while both duration groups ramped to the same threshold level, they did so with different slopes (Fig. 2D and H). Statistical analysis confirmed that there was no significant difference in the threshold signal value at the end of the waiting period across duration groups (Fig. 2F and J). However, the slopes of the ramping calcium signals differed significantly between the duration groups (2-second group: F = 6.442, p = 0.0031, n = 6 mice, one-way ANOVA; Fig. 2E; 3-second group: F = 8.421, p < 0.001, n = 6 mice; Fig. 2I). Additionally, the ramping slope showed a negative correlation with group duration (2-second group: R² = 0.375, p = 0.0015; 3-second group: R² = 0.548, p < 0.0001; n = 6 mice per group; Fig. 2E and I). These findings demonstrate that VTA DAergic neurons exhibit a characteristic ramp-to-threshold activity pattern during the waiting phase of the time perception task.

### Optogenetic manipulation of VTA DA neurons affects waiting durations

To determine whether the activity of VTA DA neuron controls timing behavior, we transiently manipulated their activity using optogenetics in 20% of pseudorandomly selected trials after the mice entered the waiting zone (Fig. 3A and B). Activating VTA DA neurons resulted in a shift of the cumulative probability distribution of waiting durations towards shorter intervals (2s group: F = 4.6, p = 0.001, n = 7 mice; 3s group: F = 5.96, p = 0.0024, n = 5 mice; one-way ANOVA; Fig. 3D to F, blue), whereas inhibition significantly shifted the distribution towards longer durations (2s group: F = 2.89, p = 0.035, n = 5 mice; 3s group: F = 6.75, p = 0.0003, n = 6 mice; Fig. 3H to J, red). Importantly, these optogenetic effects on waiting duration distributions were specific to the trials where laser manipulation occurred. Trials that did not involve laser manipulation (including those immediately preceding and following stimulation/inhibition) showed no significant differences compared to trials conducted on previous or subsequent days (p > 0.5; Fig. 3D to J). Furthermore, optogenetic manipulations did not influence running times after the mice exited the waiting zone, nor did they affect locomotion in an open field test (Fig. S3C to E). To account for non-timing factors, we conducted optogenetic manipulations in a non-timing group (0s group; immediate reward). In this group, neither activation nor inhibition of VTA DA neurons significantly altered waiting durations (Fig. S3A to D), confirming that VTA DA neurons specifically regulate timing behavior.

**Fig. 3.**
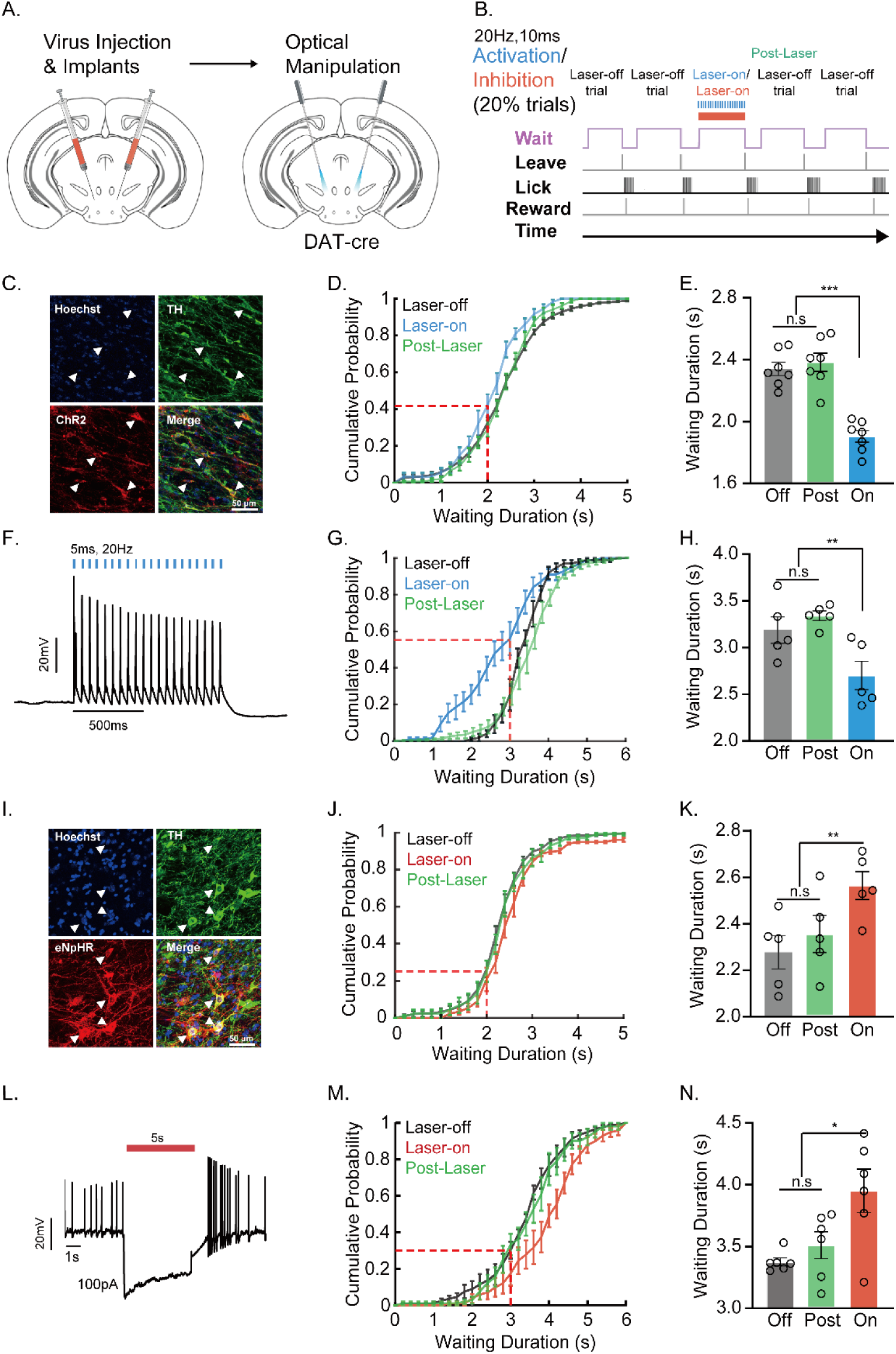
Optogenetic manipulation of VTA DA neurons bidirectionally regulates waiting duration. (A, B) Schematic of optogenetic protocol: ChR2 (activation) or eNpHR3.0 (inhibition) expressed in VTA DA neurons. Transient stimulation (blue light: 20 Hz, 10 ms pulses) or inhibition (red light: continuous 10 mW) was delivered randomly (20% trials) upon waiting zone entry. Green: post-laser epochs; Dark: laser-off epochs. (C, I) Immunostaining verifying ChR2 (C: Red: ChR2-mCherry; Green: TH; Blue: Hoechst) and eNpHR3.0 (I: Red: eNpHR3.0-EYFP; Green: TH; Blue: Hoechst) expression in VTA DA neurons. (D, G) DA neuron activation shortened waiting duration (leftward shift in cumulative distributions; 2s cohort: n=7; 3s cohort: n=5). (E, H) Mean waiting time decreased by 0.4362 s (2s) and 0.4853 s (3s) versus laser-off controls, with effects reversing in post-stimulation trials (p>0.05). (J, M) DA neuron inhibition prolonged waiting duration (rightward distribution shift; 2s: n=5; 3s: n=6). (K, N) Inhibition increased waiting by 0.2873 s (2s) and 0.5750 s (3s), attenuated in post-inhibition trials. (F, L) Electrophysiological validation: Blue light (20 Hz) evoked reliable spiking; yellow light (20 mW) suppressed activity. Statistical significance: *p<0.05, p<0.01,p<0.001,***p<0.0001.

### AHN shortened waiting duration, but Morphine did not

We investigated the effects of morphine and the dopamine reuptake inhibitor AHN 1-055 on time perception in mice. Prior to behavioral testing, we administered abdominal injections of morphine or AHN to enhance dopamine activity. The results showed that the average distribution of waiting times for the mice did not change significantly after morphine injections (D=0.197, p>0.05; Fig. 4B). Similarly, the running times of the mice did not show significant changes following morphine injections (Fig. S4C).

**Fig. 4.**
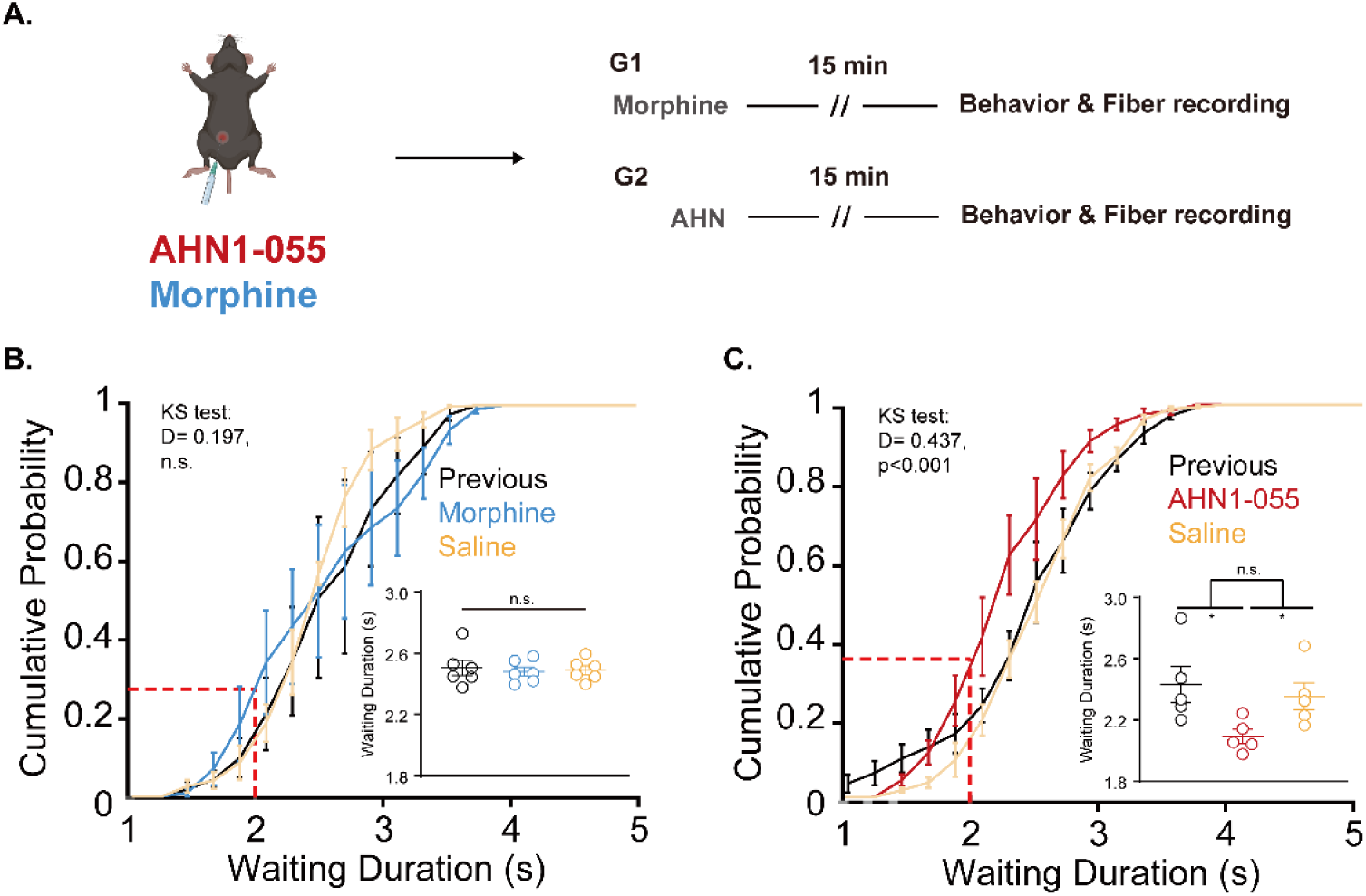
Differential effects of morphine and AHN-1055 on DA neuron calcium dynamics and waiting behavior. (A) Experimental timeline for fiber photometry recording of VTA DA neuron calcium signals following i.p. injection of morphine (1 mg/kg) or AHN-1055 (1 mg/kg). (B) Morphine (Blue) showed no significant effect on waiting duration compared to pre-injection (Light gray) or saline (Yellow) controls (mean ± SEM: 2.536±0.1, 2.516±0.1, 2.532±0.06s; n=6; Kolmogorov-Smirnov test: D=0.121, p=0.565). (C) AHN-1055 (Red) significantly shortened waiting duration, indicated by a leftward shift in cumulative distributions versus pre-injection (Light gray) and saline (Yellow) (mean ± SEM: 2.058±0.05, 2.606±0.15, 2.563±0.08s; n=5; Kolmogorov-Smirnov test: D=0.4, p<0.001). Data analyzed by two-tailed paired t-tests/ANOVA with Bonferroni correction; error bars represent SEM.

In contrast, the injection of AHN 1-055 significantly altered the mice’s waiting times compared to pre-injection levels (F=4.02, p=0.046, Fig. 4C), with five mice participating in the experiment. Specifically, the waiting time was significantly reduced after the AHN 1-055 injection (t=-2.6017, p=0.0315, Fig. 4C). The average waiting time decreased, and the probability distribution of cumulative waiting times shifted towards shorter durations after the AHN 1-055 injections (D=0.437, p<0.001; Fig. 4C), indicating that dopamine plays a role in regulating waiting times in mice.

To assess the impact of morphine and AHN 1-055 on dopaminergic neurons, we recorded the neuronal activity of these neurons in the VTA using fiber photometry. Within 10 minutes of morphine injection, calcium signals from VTA-DA neurons began to rise, while signals remained stable before the injection. By 20 minutes post-injection, the baseline calcium signal of the DA neurons was significantly elevated compared to pre-injection levels (t=4.434, p<0.01; Fig. S4A to B). In contrast, the baseline calcium signaling of VTA-DA neurons did not show significant changes after AHN injection compared to pre-injection levels (t=0.2428, p>0.05, Fig.S4D to E).

These findings suggest that systemic elevation of dopamine levels through morphine increases baseline VTA-DA neuron activity without affecting timing, whereas enhancing dopamine availability via reuptake inhibition with AHN 1-055 accelerates timing without altering baseline activity. This implies that the timing process is managed by a continuous accumulation mechanism, which compares accumulated counts with time in memory.

We hypothesized that prolonged activation of dopaminergic neurons might accelerate dopamine reuptake, consequently slowing down synaptic dopamine accumulation and extending waiting times. To test this, we optogenetically activated dopaminergic neurons during reward delivery and turned off the laser upon entry into the waiting zone (Fig. S5A). Activation resulted in longer subsequent waiting times (Kolmogorov-Smirnov test, D = 0.114, p < 0.001; Fig. S5C and E), while running times remained unchanged (D = 0.066, p = 0.52; Fig. S5D and F). This supports the notio**n** that time perception is linked to the dynamics of dopamine accumulation.

### Continuous optogenetic activation of VTA DAergic neurons prolongs post-stimulation waiting duration and reduces the slope of dopamine activity in the NAc

To further investigate the relationship between dopamine accumulation dynamics and timing, we conducted extended optogenetic activation of dopaminergic neurons while monitoring dopamine release in the nucleus accumbens (NAc) using a fluorescent dopamine sensor (Fig. 5A & Fig. S6C to D). During behavioral tasks, mice were subjected to randomized extended optogenetic stimulation of VTA dopaminergic neurons (Fig. 5B). The waiting durations measured after stimulation periods (laser off trials) were significantly prolonged, shifting the cumulative distribution toward longer values (Fig. 5C and D; Kolmogorov-Smirnov test, D = 0.271, p < 0.0001). However, running durations showed no significant differences between baseline and laser off trials (F = 0.025, p = 0.9745, n = 6 mice; Fig. 5E).

**Fig. 5.**
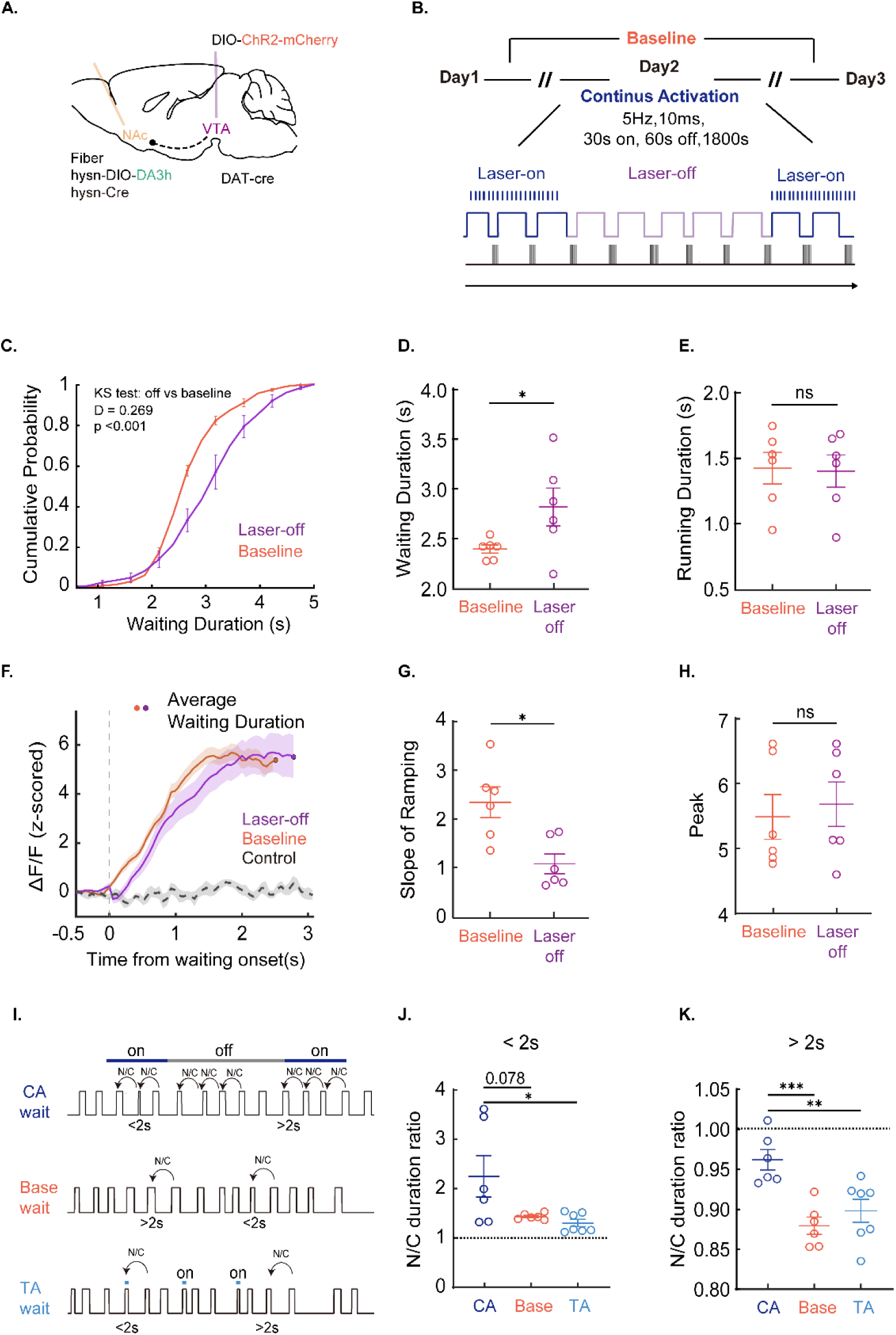
Sustained optogenetic activation of VTA DA neurons alters waiting behavior and NAc dopamine kinetics. (A) Schematic of VTA targeting (Purple: injection site; Orange: ChR2) and NAc sensor implantation (Yellow). (B) Stimulation protocol: Dark blue: 30-s light pulses (5 Hz) every 90 s on Day 2; Purple: inter-stimulation intervals; Orange: baseline days (no stimulation). (C) Cumulative distributions show prolonged waiting during laser-off intervals (Purple) versus baseline (Orange); D=0.268, p<0.0001, n=6. (D) Waiting times significantly increased during laser-off versus baseline (p<0.05). (E) Running times unchanged (n.s.). (F–H) NAc DA dynamics: aligned trajectories (F); laser-off (Purple) showed lower rise slopes despite longer waits (p<0.05, G), with unchanged peak amplitudes (n.s., H). (I–K) Trial-sequence analysis: ratios of next/current waiting time for trials <2s (J) showed no group differences (Orange: baseline; Light blue: transient activation; Dark blue: continuous activation); for trials ≥2s (K), continuous activation significantly altered ratios versus other conditions (p<0.001). Statistical tests: Kolmogorov-Smirnov and paired t-tests.

NAc dopamine signals displayed a progressive ramp during waiting, reaching the same threshold for laser-off and baseline conditions (Fig. 5F). However, following VTA dopaminergic optogenetic stimulation, the slope of the NAc dopamine ramp during the next waiting trial was significantly reduced compared to baseline condition (t = 3.826, p < 0.05; Fig. 5G), while the peak amplitude at the end of the wait remained unchanged (t = 0.3867, p = 0.7149; Fig. 5H). Subsequent regression analysis revealed a significant inverse correlation between signal slope and waiting duration (R-square = 0.098, p < 0.001; Fig. S5E), indicating slower dopamine accumulation during lase-off period. In contrast, terminal dopamine levels at the end of the wait showed no significant correlation with waiting durations (R-square = 0.0001, p = 0.9438, n = 6; Fig. S5F).

To examine how different patterns of dopamine activation influence timing behavior, we divided all individual trials into short-duration (<2 s) and long-duration (>2 s) groups based on their waiting times (Fig. 5I). We then computed the next/current ratio (N/C ratio), defined as the waiting duration of the subsequent trial divided by that of the current trial. A ratio > 1 indicates a lengthening of waiting time, whereas a ratio < 1 reflects a shortening. When the current waiting duration is shorter than 2 s (no reward delivered), mice generally increase their waiting time in the subsequent trial (ratio > 1), consistent with a negative reward prediction error (RPE) driving compensatory effort. Conversely, when the current waiting duration exceeds 2 s, mice tend to shorten their waiting time in the next trial (ratio < 1), consistent with a foraging strategy that maximizes reward rate efficiency (38–41).

For short-duration trials (<2 s), the N/C ratio was significantly higher in the continuous activation (CA) group compared with the baseline and transient activation (TA) groups (CA: ∼2.26; baseline: ∼1.41; TA: ∼1.29; F = 4.839, p < 0.05). For long-duration trials (>2 s), the CA group again showed a significantly higher N/C ratio than the baseline and TA groups (CA: ∼0.96; baseline: ∼0.88; TA: ∼0.89; t = 6, p < 0.001; Fig. 5K). These patterns are consistent with RPE and foraging theories, in which short waits induce compensatory lengthening and long waits promote shortening to optimize reward rate. Crucially, the CA group exhibited higher N/C ratios under both short- and long-duration conditions, indicating generally prolonged waiting behavior. This consistent elevation suggests that continuous dopamine activation slows dopamine accumulation dynamics, leading to longer waiting durations.

## DISCUSSION

Our study challenges the classical dopamine clock hypothesis by demonstrating that time perception is not controlled by absolute dopamine levels, but by the temporal dynamics of dopamine accumulation. Using a novel delayed-reward time perception task, we demonstrated that mice exhibit scalar timing properties consistent with Weber’s law (28, 34, 35) (Fig. 1K to M), validating the task’s ability to probe endogenous timing mechanisms. Crucially, fiber photometry signals revealed a ramping-to-threshold activity pattern in VTA dopaminergic neurons during waiting (Fig. 2). The slope of the ramping signal negatively correlates with perceived duration—slower accumulation led to longer subjective time estimates (Fig. 2E and I). This pattern suggests that dopamine acts as an “accumulator” in a neural timing circuit, with the accumulation rate determining when a temporal threshold is reached.

Numerous studies (1, 4, 6, 10, 11, 14, 15, 27) indicate dopamine activity influences time perception, often through pharmacological or genetic manipulations affecting dopamine pathways over extended periods or lifetimes. The brain’s plasticity allows it to seek balance by adjusting opposing pathways when one pathway is altered. Pharmacological experiments have shown that dopamine agonists (e.g. methamphetamine), significantly accelerate the brain’s internal clock, while dopamine receptor antagonists (e.g. haloperidol) slow it down (36). However, increasing or decreasing dopamine synthesis does not affect the internal clock (26, 37). Conditional DAT knockout speeds up timing (31), whereas overexpression of D2 receptors slows it (32). These conflicting results cannot be explained by the overall strength of dopamine activity alone. Our findings suggest that the speed of time perception depends more on the efficiency of dopamine accumulation during the perception period rather than the overall strength of dopamine activity. Both dopamine agonists and conditional DAT knockout lead to faster dopamine accumulation during timing, accelerating the clock. Conversely, overexpression of D2 receptors or the use of dopamine receptor antagonists, although appearing to enhance or weaken dopamine signaling pathways, actually increases dopamine reuptake efficiency. This slows down dopamine accumulation during time perception, thus slowing the internal clock.

Our optogenetic manipulations resolve key paradoxes. Transient activation of VTA DA neurons accelerated time estimation (shortened waiting; Fig. 3D to F), consistent with the dopamine clock hypothesis. However, prolonged activation slowed timing (Fig. 5C and D), a result incompatible with simple models based on dopamine levels. This dichotomy is explained by dopamine’s dual role: acute release directly advances the accumulator, while sustained high activity triggers compensatory mechanisms like enhanced reuptake efficiency (Fig. 5H). DAT upregulation reduces net accumulation rates per unit time, effectively decelerating the internal clock. This framework accounts for why: (i) DAT knockout (increasing accumulation) speeds timing; (ii) D2 receptor overexpression (increasing reuptake) slows timing; (iii) systemic morphine (elevating baseline without altering reuptake dynamics) does not affect timing (Fig. 4B); and (iv) reuptake inhibition (AHN 1-055, boosting accumulation rates) shortens waiting time (Fig. 4C).

We propose a “Dynamic Dopamine Accumulation Threshold” (DDAT) model comprising three key components: (i) VTA DA neurons ramp up activity at a slope inversely proportional to target duration; (ii) accumulated dopamine reaching a fixed threshold triggers action initiation; (iii) chronic DA elevation induces DAT plasticity, modulating the accumulation slope. This model explains why optogenetic inhibition prolongs waiting (reduced slope; Fig. 3H to J) and acute activation shortens it (steeper slope; Fig. 3D to F). Crucially, it reconciles disparate pharmacological and genetic findings by emphasizing temporal dynamics over static levels.

Dopamine is also well known for conveying reward prediction errors (RPEs) (38–41), thereby driving adaptive adjustments in subsequent behavioral strategies. In our data, following short waits (<2 s), all conditions (laser on/off) showed adaptive lengthening of subsequent waits (ratio > 1; Fig. 5J), consistent with the notion that a negative RPE signals an “under-rewarded” state that promotes compensatory effort. After long waits (>2 s), all conditions exhibited adaptive shortening (ratio < 1; Fig. 5K), reflecting a foraging strategy that optimizes reward rate. Critically, the ratios under prolonged optogenetic activation were significantly higher after both short and long waits, indicating slower dopamine accumulation—consistent with the dynamics observed in laser-off trials during prolonged activation (Fig. 5F to H). To account for this slowed accumulation under prolonged stimulation, we propose that sustained optogenetic activation enhances dopamine reuptake efficiency, thereby reducing the net rate of dopamine build-up and consequently lengthening waiting durations

By demonstrating that the dynamics of dopamine accumulation, rather than its absolute levels, control time perception, our work overturns the dopamine clock hypothesis. The DDAT model unifies decades of conflicting evidence, positioning dopamine reuptake efficiency as a critical regulator of internal timing. This paradigm shift opens new avenues for treating timing disorders and prompts a reinterpretation of dopamine’s role in decision-making and reward processing.

Contrary to the view that instantaneous dopamine concentration directly sets an internal clock, our waiting-based paradigm demonstrates that integrated, cumulative dopamine dynamics—shaped by release and reuptake—causally determine temporal judgments. This distinction has practical implications for artificial humoral–neural networks and neuromorphic architectures: implementing modulatory channels that accumulate and decay over extended timescales (rather than acting as transient, instantaneous gain changes) should enable more stable and flexible timekeeping, improve robustness to noise, and reconcile local versus global modulatory effects in optimization and decision-making tasks.

## MATERIALS AND METHODS

### Mice

DAT-Cre mice (006660), C57BL/6J mice were purchased from the Jackson Laboratory. Virus validation experiments used 4-week-old mice, while other experiments used 6- to 8-week-old mice with a body weight of 19-25g. Mice were group-housed and maintained on a 12 h light-dark cycle (i.e., light cycle; 9 am–9 pm) with food and water available ad libitum. The holding rooms had a constant temperature of 24°C, a humidity of 46%, a pressure of 9.9 Pa, and free access to food and water except during behaviour training. The animals were acclimatized for one week before the start of the experiment, with a daily 5-minute routine of handling the animals at a scheduled time. All experimental manipulations complied with the ethical guidelines for experimental animals of Peking University and the Chinese Institute for Brain Research and adhered to the regulations issued by the national and the Institute’s Animal Experimentation Centre.

### Preparation of adeno-associated viruses (AAVs)

Adeno-associated virus (AAV) with a titer exceeding 2 × 10^12 vector genomes (vg) mL−1 was utilized in all experiments. The rAAV9-hSyn-DIO-DA3h (BC-0859) was procured from BrainCase Co., Ltd. The following constructs were obtained from BrainVTA Co., Ltd.: rAAV9-hSyn-DIO-hM3D (Gq)-mCherry (PT-0019), rAAV9-EF1α-DIO-hChR2-mCherry (PT-0002), and rAAV9-EF1α-DIO-eNpHR3.0-mCherry (PT-0006). Additionally, rAAV9-EF1α-DIO-GCaMP6m and rAAV9-hSyn-Cre were produced and packaged by the vector core of the Chinese Institute for Brain Research. All viral preparations were stored in aliquots at −80°C until needed.

For the neuronal population recording experiment, AAV9-EF1α-DIO-GCaMP6m and AAV9-EF1α-DIO-EGFP were administered to DAT-Cre mice. In the optogenetic stimulation experiment targeting ventral tegmental area (VTA) dopaminergic neurons, AAV9-EF1α-DIO-hChR2(H314R)-mCherry was employed for activation, AAV9-EF1α-DIO-eNpHR3.0-mCherry was utilized for inhibition, and AAV9-EF1α-DIO-mCherry served as a control to mitigate the effects of light on behavior. For the experiment assessing dopamine release during the waiting period, rAAV9-EF1α-DIO-hChR2-mCherry was administered to VTA neurons for activation, while rAAV9-hSyn-DIO-DA3h and rAAV9-hSyn-Cre were co-injected into the nucleus accumbens (NAc) of DAT-Cre mice to monitor dopamine dynamics in the NAc.

### Stereotaxic viral injection and optical fiber implantation

Six-week-old male mice were anesthetized using isoflurane (3.5% induction, 1.5-2% maintenance). Once deep anesthesia was established, mice were positioned on a stereotaxic instrument (Stoelting Co., USA). Anesthesia was maintained at 1% to 1.5% isoflurane delivered through an anesthesia nosepiece. The eyes were coated with erythromycin eye ointment. The scalp was incised, and the fascia over the skull was removed using a solution of 3% hydrogen peroxide in saline. The bregma and lambda points were used to ensure the mouse head was properly leveled.

For the neuronal population recording experiment, a 200 to 300 μm window was drilled above the VTA (coordinates: AP = −3.20 mm, ML = 0.25 mm, DV = −4.3 mm) for the viral injection and optical fiber implantation (coordinates for fiber: AP = −3.20 mm, ML = 0.25 mm, DV = −4.15 mm). A total of 100 nl of AAV solution was slowly injected unilaterally at a rate of 30 nl/min (this speed was used in all the following experiments) for fiber photometry recording, using an MO10 device (Narishige, Japan) coupled with a glass electrode. A total of 160 nl of AAV solution was slowly injected bilaterally (80 nl at each injection site) for optogenetic stimulation. A total of 100 nl of AAV solution was slowly injected unilaterally to facilitate the recording of dopamine dynamics in the NAc.

The coordinates for optogenetic stimulation of DAergic neurons in the VTA are AP = −3.20 mm, ML = ±0.99 mm, DV = −4.19 mm, with an angle of 10 degrees. The coordinates for recording dopamine dynamics during periods of waiting in the nucleus accumbens (NAc) are AP = 0.75 mm, ML = 1.00 mm, DV = −4.65 mm. Following the injection, the glass electrode was held in place for 10 minutes before being slowly withdrawn. An optical fiber (200 μm, 0.37 numerical aperture, Originopto, Hangzhou) housed in a ceramic ferrule was then carefully inserted into the brain tissue, with the tip positioned 150 μm above the viral injection site. The fiber was secured to the skull with dental cement. Postoperative care included maintaining normothermia during the recovery from anesthesia and administering subcutaneous enrofloxacin prophylaxis (Baytril®; 10 mg/kg, twice daily) for 72 hours. After a 14-day recovery period with daily veterinary monitoring, the animals underwent either VTA neuronal population recording assessments, optogenetic stimulation experiments, or dopamine dynamic recording together with optogenetic stimulation and behavior tasks.

### Fiber photometry recording and calcium data analysis

#### Fiber recording

DAergic neuronal activities in the VTA and dopamine dynamics in the NAc during the waiting task were monitored using an FPS-405/470/578 photometry system developed by the CIBR Imaging Core, along with the Tri-photometry software. We employed a dual-wavelength approach, utilizing a 470 nm LED to stimulate calcium signal-dependent fluorescence, while a 405 nm LED reference channel provided motion correction. The light intensity at the fiber tip was calibrated to 10–20 μW. We alternated excitation wavelengths and collected fluorescence signals using a CMOS camera at a rate of 10 frames per second for each channel during dual-color imaging. Three tag wires were extended from the data acquisition card of the fiber photometry system and connected to the pins of a microcontroller. Through interaction between the microcontroller and MATLAB, TTL signals were generated as high-level voltage pulses during specific behavioral events—such as entry into the waiting zone, licking behavior, and water delivery, the latter also triggered by the microcontroller. These TTL pulses were then transmitted back to the event recording channel of the fiber photometry system via the microcontroller. This method was primarily implemented to synchronize behavioral video with calcium signals for subsequent analysis. Additionally, we used OBS Studio (https://github.com/obsproject/obs-studio) to record and save each fiber photometry session.

To minimize autofluorescence of the optical fiber, the recording fiber was photobleached for 24 hours using a high-power LED before recording. The photometry data were analyzed using a custom-written MATLAB (MATLAB R2023b, MathWorks) program, and background autofluorescence was subtracted from the recorded signals. The final signal was corrected by motion and bleaching using MATLAB codes.

#### Calcium data analysis

Each raw fluorescence signal was exported and analyzed using MATLAB (R2023b, MathWorks) with custom scripts. Initially, photo-bleaching corrections were applied to each channel. The signal from 405 nm channel was subsequently utilized to correct for motion in the calcium signal channel (41). Event channels, including waiting, licking, and reward, were automatically filtered using MATLAB’s differential function to identify the onset and offset times for each event. The duration of waiting was calculated by subtracting the waiting offset time from the waiting onset time, while the running duration was determined by subtracting the initial valid lick from the corresponding waiting offset.

We extracted the fluorescence data (F) for each waiting period, defined as 0.5 seconds before the waiting onset to 3 seconds after the waiting onset for the 2-second group, and to 4 seconds after the waiting onset for the 3-second group. This data was then transformed into z-scores using the following formula:

F_zscore=(F-mean(F_pre)/s.d.(F_pre)

where Fpre is defined as the fluorescence signal 0.5 seconds prior to waiting onset. The change in the calcium signal was calculated by subtracting the peak value of the Fzscore signal. The signal traces are presented as mean plots, with a shaded area representing the standard error of the mean (SEM) of fluctuations.

### Electrophysiology recording

In the validation experiments for opsin-expressing viral efficacy, whole-cell patch-clamp recordings were performed to assess action potential (AP) firing in VTA neurons expressing ChR2-mCherry or eNpHR3.0-mCherry under blue (470 nm) or orange (590 nm) light stimulation, respectively. The experimental protocol was as follows:

rAAV2/9-EF1α-DIO-hChR2-mCherry or rAAV2/9-EF1α-DIO-eNpHR3.0-mCherry was stereotaxically delivered into the VTA of DAT-Cre transgenic mice to target dopaminergic neurons. Two weeks post-injection, acute brain slices were prepared for electrophysiological recordings. Approximately 100 mL of ice-cold slicing solution (pre-frozen at −80°C for 30 min) was manually fragmented into a slush-like consistency, oxygenated with carbogen (5% CO₂/95% O₂), and transferred to a slicing chamber. Mice (4–6 weeks old) were anesthetized with isoflurane, followed by rapid brain extraction and trimming. The tissue block was secured to a vibratome stage (VT1200S, Leica) using cyanoacrylate adhesive and submerged in the chilled slicing solution.

Coronal midbrain slices (300 μm thickness) containing the VTA were sectioned at 0.8 Hz oscillation frequency and 0.8 mm/s advance speed. Slices were incubated in oxygenated artificial cerebrospinal fluid (ACSF) at 34°C for 30 min, then maintained at room temperature (24–26°C) for ≥30 min before recording. During experiments, slices were continuously perfused with oxygenated ACSF (2 mL/min) at room temperature.

Patch pipettes (borosilicate glass: 1.0 mm OD, 0.86 mm ID; P-97 puller, Sutter Instrument) with 4–6 MΩ resistance were used for whole-cell current-clamp recordings. Signals were amplified (Multiclamp 700B, Axon Instruments), digitized (Digidata 1500B), and analyzed using Clampfit 10 (Molecular Devices) and Mini Analysis Program (Synaptosoft). Optical stimulation (Polygon 400, Mightex) consisted of 470 nm blue light (20 Hz, 5 ms pulses) or 590 nm orange light (5 s duration).

ChR2-expressing neurons reliably followed 20 Hz blue light stimulation with phase-locked APs (Fig.3a-b), confirming effective optogenetic activation. In eNpHR3.0-expressing neurons, orange light illumination during 100 pA depolarization-induced firing completely suppressed action potentials, with full recovery post-illumination, demonstrating reversible inhibition of VTA-DA neuronal activity.

### Behavioral tasks

Two weeks after the fiber photometry cannula implantation surgery, mice were placed on a water restriction protocol. Their body weights were monitored daily, and water access was controlled to maintain them at 85–90% of their baseline free-drinking body weight for five consecutive days. This water restriction period was critical for enhancing the mice’s motivation to obtain water rewards, ensuring high levels of engagement in the upcoming task. Subsequently, the experimenter performed gentle handling for 5 minutes per day over three consecutive days to habituate the mice to human interaction. Task training commenced immediately after this handling period. All behavioral tasks were conducted during the dark (active) phase of the light/dark cycle under dim red light.

The foraging task utilized a shuttle box comprising two chambers (10 cm × 10 cm × 15 cm) connected by a corridor (45 cm × 5 cm × 15 cm; see Fig. 1a-b). A water port (1.2-mm O.D. steel tube, positioned 3 cm above the floor) was installed at one end of the chamber, designated as the reward zone, while the opposite chamber served as the waiting zone. The position of the mouse within the shuttle box was monitored in real-time using a custom MATLAB (2023b, MathWorks) program and an overhead camera (ZhongAoWeiKe, PCBA-P1080P). Experimental procedure control and behavioral event data acquisition were facilitated through a custom MATLAB program and an integrated circuit board (Arduino UNO R3).

#### Pre-training

A water-restricted mouse was placed in the shuttle box for free exploration for up to 1 hour. When the animal traveled from the waiting zone through the corridor to the reward zone to lick the water port, 10 μl of water was delivered by a step motor in 100 ms as a reward. A capacitor sensor monitored the timing and duration of licking. The animal returned to the waiting zone to initiate the next trial. The time spent in the waiting zone was defined as the waiting duration. The training was conducted every day for a week. All mice learned to move quickly back and forth between the two chambers to maximize the reward rate within 1 week.

#### Time-waiting-task

From the second week, the volume of water reward was changed to a settled rule: a wait time of 0 to 2 s for 0 μl; above 2s triggered delivery of 10 μl (2s training condition); and above 3s triggered delivery of 10 μl (3s training condition) as shown in Fig. 1A. The training was conducted 7 days a week, from Monday to Sunday.

### Open-field test in mice

For the control experiment, the locomotion of DAT-cre mice was tested in the Open-field behaviour assay for two conditions: optogenetical activation and inhibition. The open-field test was performed in ENV-510 test chambers (27.3 × 27.3 × 20.3 cm3, Med Associates). The chamber is equipped with infrared photo beams, which were used to evaluate spontaneous mouse locomotor activity. At the start of the test, mice were placed in the center of the arena, and were allowed to freely explore the environment for 10 min. Mouse movement was tracked and analyzed by the Activity Monitor 7 software (Med Associates) for the total travel distance, the number of entries to the central zone (14.29 × 14.29 cm2) and the time spent in the central zone.

### Optogenetic stimulation

Dual wavelength optogenetic stimulation protocols were implemented using 488 nm (activation) and 589 nm (inhibition) lasers following established methodologies.

#### Transient Stimulation Protocol

After 3 weeks of time-restricted waiting task training, animals received pseudorandom optical interventions during 20% of test trials. Activation parameters consisted of 10 Hz square-wave pulses (20 ms width), while inhibitory protocols employed continuous illumination. Stimulus delivery was contingent on behavioral positioning: laser activation initiated upon entry to the waiting zone and ceased upon exit, with a maximum exposure duration cap of 16 s regardless of prolonged zone occupancy. Terminal optical power measured 10.0 ±0.5 mW (488 nm) and 8.0 ±0.5 mW (589 nm) at the bifurcated fiber tip.

#### Continuous Stimulation Protocol

Optogenetic modulation commenced 200 s post-task initiation. Activation cycles (5 Hz, 10 ms pulse width) followed an intermittent pattern: 30s ON / 60s OFF intervals repeated across an 1,800s experimental window. All stimulation sequences were manually discontinued if behavioral sessions exceeded protocol duration limits. Power calibration matched transient protocol specifications.

### Drug preparation and injection

For morphine administration, animals received intraperitoneal (i.p.) injections of morphine (1 mg/kg), prepared as a drug solution dissolved in 0.9% sodium chloride, utilizing sterile 1 mL syringes. AHN 055 (MedChemExpress) was administered in a similar manner at a dose of 1 mg/kg, prepared in phosphate-buffered saline and delivered via the same route. Injections were conducted under brief restraint, and all drug solutions were freshly prepared prior to each experimental session. Following pharmacological treatment, subjects were transferred to the temporal chamber for monitoring. After a 15-minute post-injection interval, which allowed systemic drug distribution and onset of action, each mouse was carefully placed in the waiting zone to initiate the time-waiting task.

### Immunohistochemistry

Mice were anaesthetized (using Avertin) and intracardially perfused with PBS followed by 4% PFA in PBS buffer, and brains were dissected and fixed at 4 °C overnight by 4% PFA in PBS. Brains were sectioned at 40-μm thickness using a VT1200 vibratome (Leica). Slices were placed in blocking solution containing 5% normal goat serum, 0.1% Triton X-100 and 2 mM MgCl2 in PBS for 1 h at room temperature. For the co-localization experiments, slices were incubated in AGT solution (0.5% normal goat serum, 0.1% Triton and 2 mM MgCl2 in PBS) containing primary antibodies, anti-TH antibody (Abcam, catalog no. ab13970, chicken, dilution 1:1,000) and anti-c-Fos antibody (Rockland, catalog no. 600-401-379, rabbit, dilution 1:500), overnight at 4 °C. On the following day, slices were rinsed three times in AGT and incubated for 2 h at room temperature with secondary antibodies: Goat anti-Chicken Alexa Fluor 488 (Abcam, catalog no. ab150169, dilution 1:500) and Goat anti-Rabbit iFluor 555 (AAT Bioquest, catalog no. 16690, dilution 1:500). After three washes in AGT, slices were incubated in AGT containing DAPI (MedChemExpress, catalog no. HY-D0814, 5 mg ml−1, dilution 1:1,000) for 15 min at room temperature and then rinsed in PBS. Slices were mounted on slides. Slices were imaged on an Aperio Versa (Leica) under a ×10 objective and ×40 objective for confocal imaging.

### Slope analysis

Fiber photometry datasets were preprocessed through a custom MATLAB GUI adapted from previously described methodologies. Z-score normalized ΔF/F signals underwent trial-wise segmentation and subsequent reanalysis via dedicated MATLAB scripts. Trials were computationally sorted by descending waiting duration, with photometric traces truncated to exclude signal segments exceeding trial-specific waiting durations (excess values designated as missing data, NaN). Linear regression analysis was performed on individual trial ΔF/F trajectories to derive activation slope parameters, which were then paired with corresponding behavioral wait times. Statistical inclusion criteria required positive regression coefficients (slope>0) meeting stringent significance thresholds (p<0.001, uncorrected). Qualifying slope values were subsequently analyzed through linear regression against their associated waiting durations to establish behavior-neural dynamic correlations.

### Quantification and statistical analysis

Except where indicated otherwise, all summary data are presented as the mean ± s.e.m. Imaging data were processed using ImageJ (1.53c) or MATLAB software (matlab R2020a and R2022a) and plotted using OriginPro 2020b (OriginLab), GraphPad Prism 8.0.2 or Adobe Illustrator CC. The SNR was calculated as the peak response divided by the standard deviation of the baseline fluorescence. Group differences were analyzed using a one-way ANOVA with Tukey’s multiple comparison test, a one-way ANOVA with Dunnett’s multiple comparison test, a two-way ANOVA with Sidak’s multiple comparison test, a two-tailed Student’s t-test or a mixed-effects model (GraphPad Prism 10.0.2). Differences were considered significant at p<0.05; *p<0.05, **p<0.01, ***p<0.001, ***p<0.0001 and not significant (p>0.05). For all representative images and traces, similar results were obtained for >3 independent experiments.

## ACKNOWLEDGEMENTS

This work was supported by China STI2030-Major Projects 2021ZD0202800 to W.S. The National Natural Science Foundation of China (32200837) grant to Z.-L.G. The Beijing Postdoctoral Research Program (No. ZZ-2025-75) to T.L. We thank M.S. Xinwei Gao and Dr. Qingchun Guo from the CIBR Imaging facility for microscopic imaging, Dr. Fei Zhao from the CIBR vector core for AAV packaging, and all members of the Sun Lab and other colleagues from CIBR for their feedback on this work.

## Author contributions

Conceptualization: W.S., Z.G., T.L. Methodology: W.S., Z.G., T.L., J.S., Y.L. Investigation: T.L., J.S., Y.L., A.L. Funding acquisition: W.S., Z.G., T.L. Supervision: W.S., Z.G. Writing – original draft: W.S., Z.G., T.L. Writing – review & editing: W.S., Z.G., T.L.

## DATA AVAILABILITY

All data are available in the main text or the supplementary materials. Further details can be obtained from the corresponding author upon request.

## SUPPLEMENTARY MATERIALS

### Supplementary Figures

**Fig. S1.**
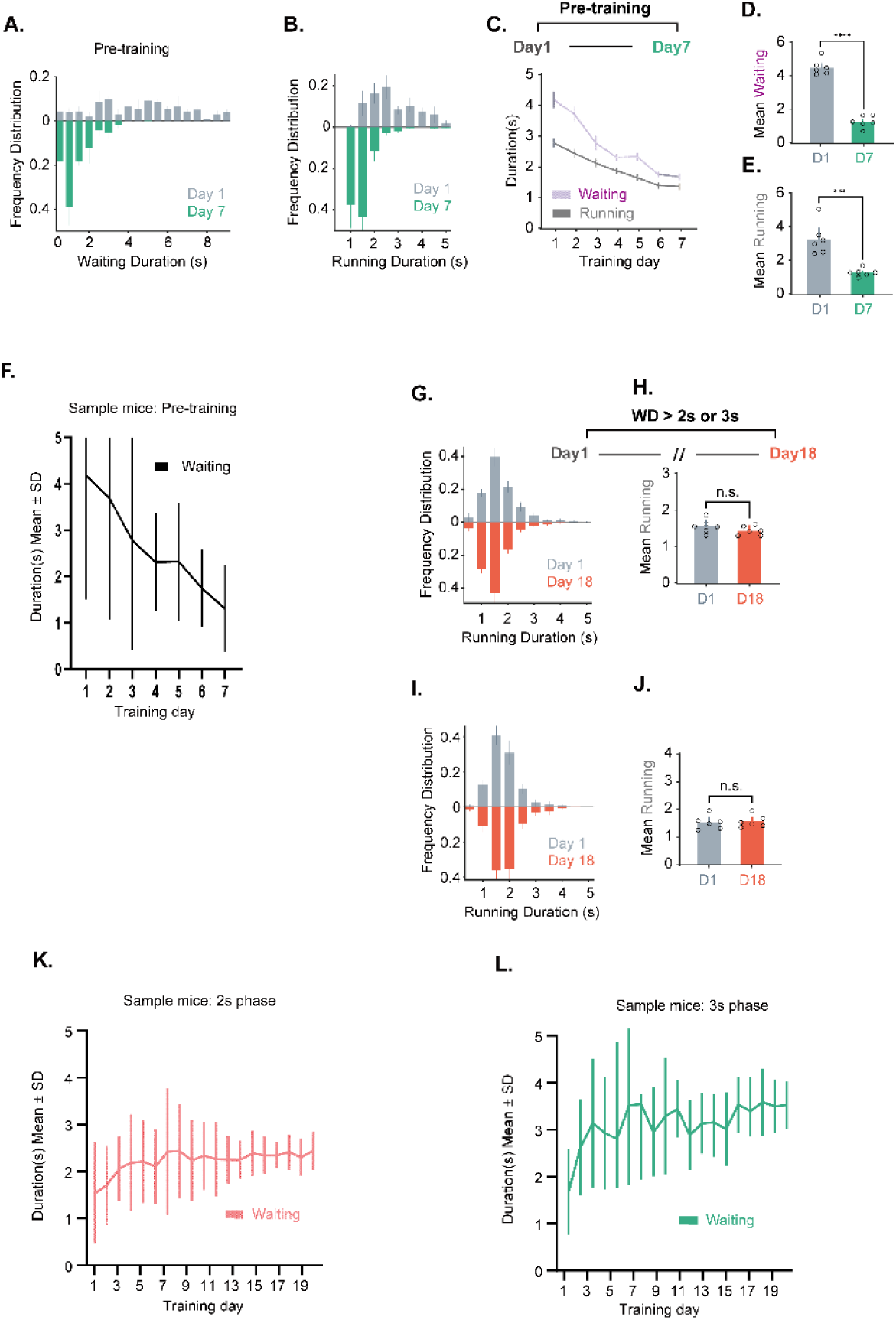
Pre-training outcomes and individual mouse performance samples. (A–E) Behavioral results during pre-training: Days 1–7: pre-training phase. (A) Waiting duration distributions on Day 1 (gray) were broadly distributed, whereas Day 7 (green) showed tight clustering near 1 s. (B) Movement durations stabilized by Day 7 (green), indicating mastery of reward acquisition. Gray: Day 1; Green: Day 7. (C–E) Across pre-training: Waiting duration significantly decreased from ∼4.2 s to ∼1.8 s (purple). Movement duration decreased from ∼3 s to ∼1.5 s (gray; n = 6). (F) Representative pre-training trajectory of an individual mouse. (G–J) Movement duration analysis for 2-second and 3-second cohorts: formal training for waiting >2 s (or >3 s in a separate cohort) commenced on one day after pre-training. Formal training days are represented from day 1 (gray) to day18 (coral). (G) Day 1 vs. Day 18 movement duration distributions (2-second cohort). (H) No significant difference in 2-second cohort movement durations (paired t-test). (I) Day 1 vs. Day 18 movement duration distributions (3-second cohort). (J) No significant difference in 3-second cohort movement durations. (K–L) Exemplar training trajectories: (K) 2-second cohort mouse: Mean waiting duration ± SD (red dashed line: 2 s target). (L) 3-second cohort mouse: Mean waiting duration ± SD (red dashed line: 3 s target).

**Fig. S2.**
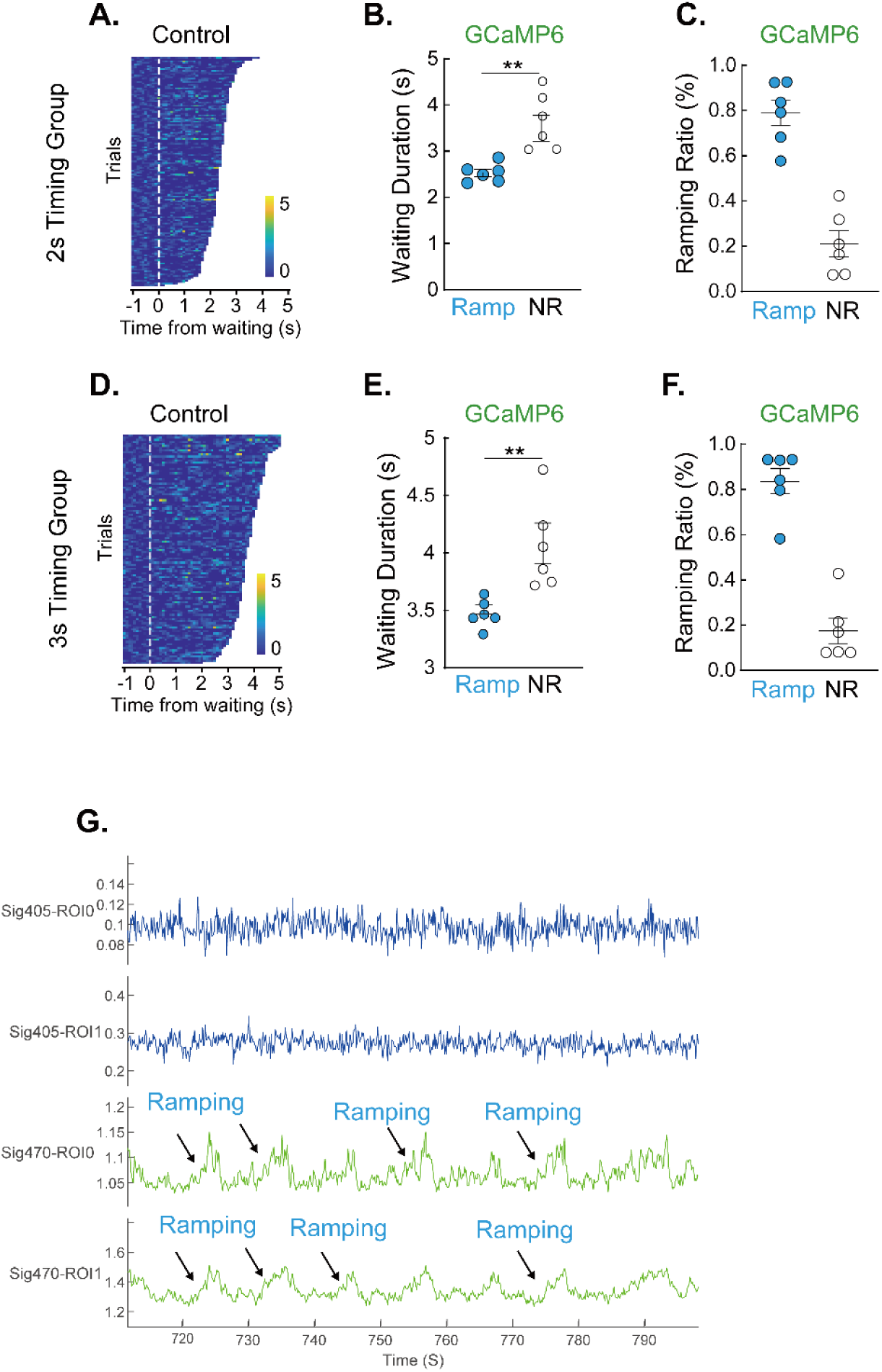
Ramping activity in VTA-DA neurons during 2-second and 3-second waiting trials, and heatmap of GFP controls for the same training protocol. (A) Heatmaps sorted by waiting duration (representative mouse) for 2s mice injected with AAV-GFP as a control. (B) Scatter plots compare waiting durations between ramping (blue) vs. non-ramping (gray) trials (2-second: n = 6; paired t-test). (C) Scatter plots compare the ratio of ramping (blue) vs. non-ramping (gray) trials (2-second: n = 6; paired t-test). (D) Heatmaps sorted by waiting duration (representative mouse) for 2s mice injected with AAV-GFP as a control. (E) Scatter plots compare waiting durations between ramping (blue) vs. non-ramping (gray) trials (3-second: n = 6; paired t-test). (F) Scatter plots compare the ratio of ramping (blue) vs. non-ramping (gray) trials (3-second: n = 6; paired t-test). (G) A demographic figure shows the raw trace of mice performing the waiting task. The neuronal activity of VTA DAergic neurons ramp up while waiting and reach its peak at reward onset. We did not observe neuronal activity in the 405 channels, indicating that the ramping activity showed up in the 470 channels came from green fluorescent proteins expressed on the DAergic neurons. Blue: the recording of 405 channels in the same mice performing the waiting task. Green: the recording of DAergic neuron activity using a *cre*-dependent Calcium indicator.

**Fig. S3.**
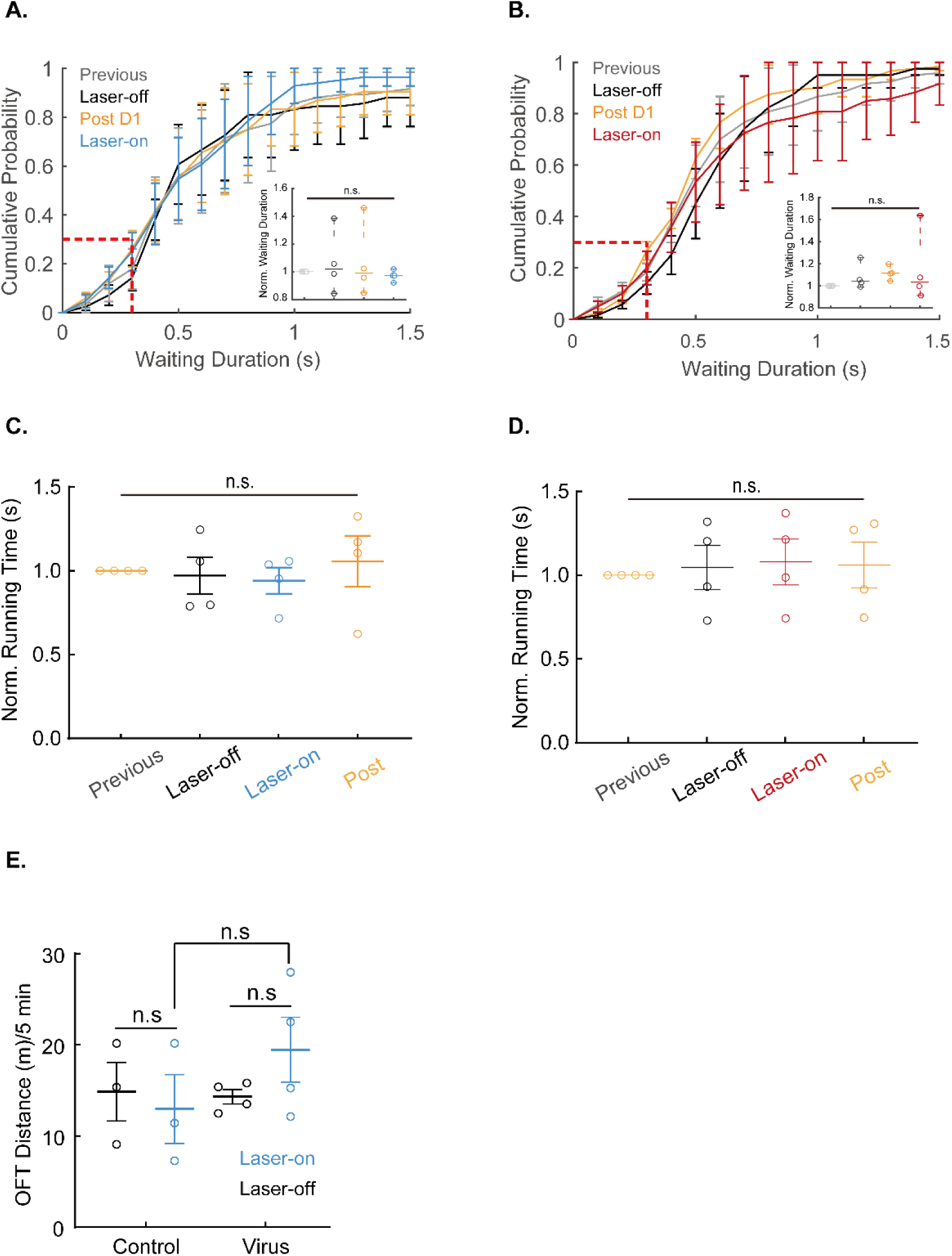
Optogenetic stimulation of VTA DA neurons does not significantly alter locomotor activity in the open field test, nor did it significantly change waiting duration in the pre-training mice. (A-B) In the pre-training group (immediate reward paradigm): Neither activation (blue, 20 Hz blue light) nor inhibition (red, continuous yellow light) of VTA DAergic neurons altered waiting duration distributions (Kolmogorov-Smirnov test) or mean waiting time (paired t-test/ANOVA, n=4). Gray (previous), Dark (no light), orange (post-stimulation day), blue (activation), red (inhibition). (C-D) Neither activation (blue, 20 Hz blue light) nor inhibition (red, continuous yellow light) of VTA DAergic neurons altered mean waiting time (paired t-test/ANOVA, n=4). Gray (previous), Dark (no light), orange (post-stimulation day), blue (activation), red (inhibition). (E) Intermittent activation of VTA DA neurons (20 Hz, 10 ms blue light pulses every 300 s) showed no significant effect on total travel distance compared to mCherry controls (n=4 mice/group). Control: AAV-DIO-mCherry, n=3; Virus: AAV-DIO-ChR2-mCherry, n=4. These experiments confirm that VTA DA neuron stimulation does not produce confounding effects on motivation, locomotor capacity, or reward perception under baseline conditions.

**Fig. S4.**
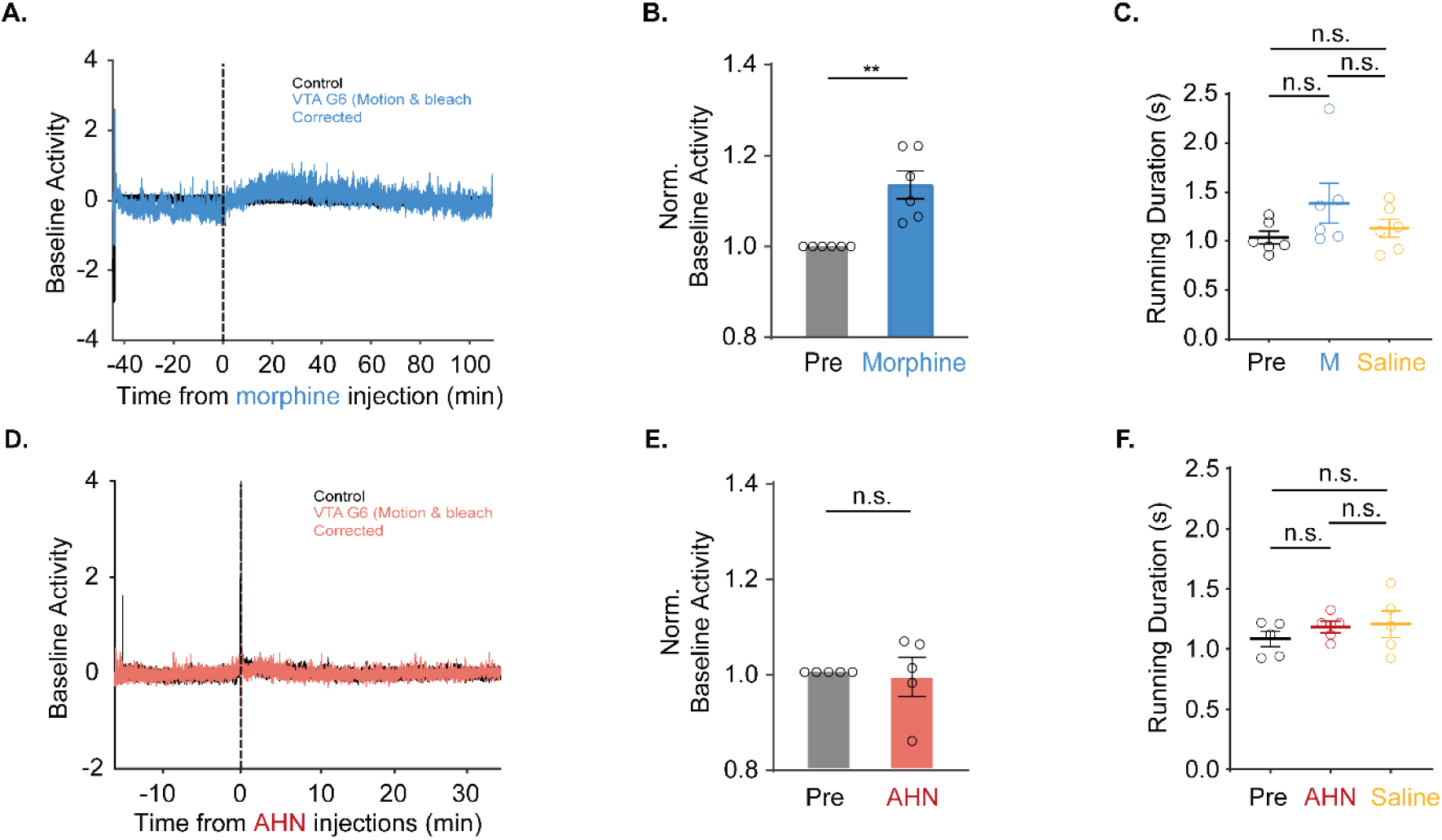
Pharmacological modulation of VTA DA neuron calcium dynamics differentially affect baseline activity and signal escalation. (A) A demographic figure shows the effect of morphine injection on DA neuronal GCaMP responses. (B) normalized ΔF/F of baseline activity significantly increased from 1 to 1.04 ± 0.01, **p<0.01, paired test). (C) Morphine (1 mg/kg i.p., blue) did not significantly reduce running time vs baseline (gray, p=0.19, paired t-test, n=6). (D) A demographic figure shows the effect of AHN1-055 (a dopamine reuptake inhibitor) injection on DA neuronal GCaMP responses. (E) There was no significant difference in normalized ΔF/F (range 0.85-1.05) for pre-injection and AHN group. (F) AHN-1055 (1 mg/kg i.p., red) showed no locomotor effects (p=0.32, n=5). Color code: Gray (pre-injection), Blue/Red (post-drug; blue: morphine, red: AHN1-055), Yellow (saline). Statistical reporting: Mean ± SEM. All comparisons verified for shuttle time consistency (p>0.1 between groups).

**Fig. S5.**
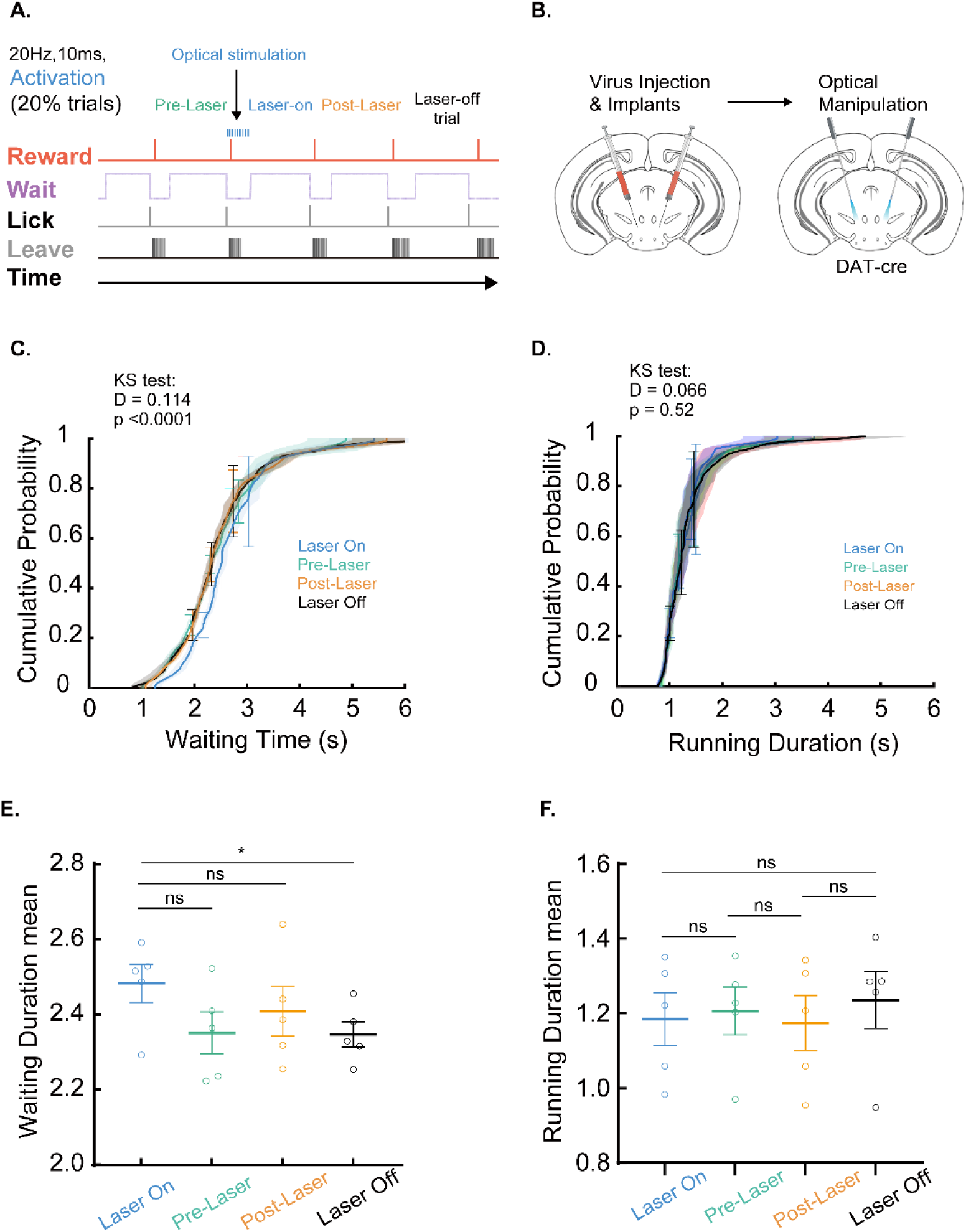
Post-reward activation of DA neurons prolongs subsequent waiting duration without affecting locomotion. (A) Experimental design. Optogenetic stimulation (20 Hz, 5-ms blue light pulses) was delivered during reward consumption in 20% randomly selected trials. Laser-on trials (blue) represent waiting periods immediately following optical stimulation applied during reward consumption. Pre-stimulation (green) and post-stimulation (yellow) trials were analyzed against non-stimulated controls (gray). (B) Injection site for optogenetic virus and fiber implantation location. (C) Cumulative waiting duration distributions for laser-on trials showed significant rightward shift versus laser-off (D=0.114, p<0.0001, Kolmogorov-Smirnov test, n=5 mice). No significant shifts occurred between pre-laser, post-laser, and laser-off conditions. (D) Running duration distributions did not significantly differ between laser-on and laser-off trials (D=0.066, p=0.52). Laser on (blue) indicates waiting trials immediately after transient DA activation during reward consumption. Pre-laser indicates waiting trials before laser-on without stimulation. Post-laser indicates waiting trials after laser-on without stimulation. Laser-off includes all non-stimulated trials (pre-laser and post-laser). (E) Mean waiting durations were laser-on 2.483±0.1136, pre-laser 2.351±0.1254, post-laser 2.408±0.1476, and laser-off 2.347±0.0755. Laser-on trials showed prolonged waiting durations with borderline significance (One-way ANOVA F=1.439, p=0.2684; laser-on vs laser-off paired t-test p<0.05, n=5). (F) Mean running durations were 1.184±0.1584, pre-laser 1.206±0.1439, post-laser 1.174±0.1651, and laser-off 1.236±0.1707. Running durations remained unchanged (One-way ANOVA p=0.24, n=5).

**Fig. S6.**
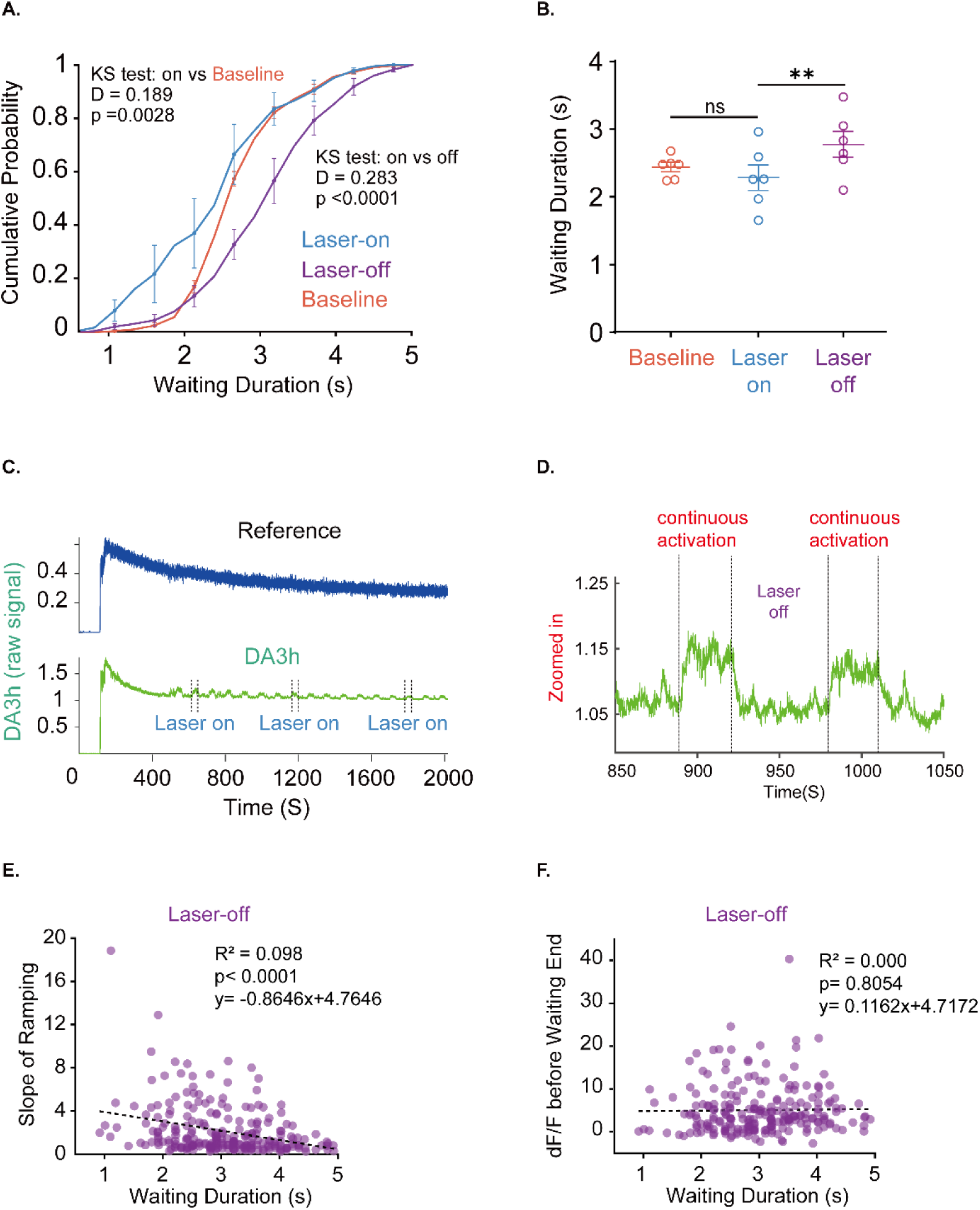
Dynamic temporal modulation of waiting behavior by sustained optogenetic activation. (A) Cumulative probability distribution of waiting durations shifted significantly leftward during sustained DA activation (laser on) versus baseline, and laser off conditions (on vs off, D=0.283, p<0.0001; on vs baseline, D=0.189, p<0.01, Kolmogorov-Smirnov test, n=6 mice). Blue indicates laser on, purple indicates laser off, orange indicates baseline. (B) Mean waiting durations for baseline (2.4742±0.1298), laser on (2.284±0.4578), and laser off (2.792±0.4664) groups (n=6). Laser on and laser off groups differed significantly (paired t-test, p<0.01). Color coding matches (A). (C-D) Dopamine photometry validation. DA transients were observed during light-on epochs (black bars) and were absent in 405nm isosbestic control. The amplitude of dopamine levels was quickly improved when VTA dopaminergic neurons were ontogenetically activated (D) and decreased after light stimulation was aborted. (E) Scatter plot and linear regression of ramping slope versus waiting durations for laser off group. Significant negative linear relationship (p<0.0001, n=6). (F) Scatter plot and linear regression of peak DA activity versus waiting durations for lase off group. No significant linear relationship was observed between the DA activity before waiting end and waiting durations (p=0.8054, n=6).

